# An automated statistical technique for counting distinct multiple sclerosis lesions can recover aspects of lesion history and provide relevant disease information

**DOI:** 10.1101/212860

**Authors:** Jordan D. Dworkin, Kristin A. Linn, Ipek Oguz, Greg M. Fleishman, Rohit Bakshi, Govind Nair, Peter A. Calabresi, Roland G. Henry, Jiwon Oh, Nico Papinutto, Daniel Pelletier, William Rooney, William Stern, Nancy L. Sicotte, Daniel S. Reich, Russell T. Shinohara, the NAIMS Cooperative

## Abstract

**Background:** Lesion load is a common biomarker in multiple sclerosis, yet it has historically shown modest associations with clinical outcomes. Lesion count, which encapsulates the natural history of lesion formation and is thought to provide complementary information, is difficult to assess in patients with confluent (i.e. spatially overlapping) lesions. We introduce a statistical technique for cross-sectionally counting pathologically distinct lesions.

**Methods:** MRI is used to assess the probability of lesion at each location. The texture of this map is quantified using a novel technique, and clusters resembling the center of a lesion are counted.

**Results:** Validity was demonstrated by comparing the proposed count to a gold-standard count in 60 subjects observed longitudinally. The counts were highly correlated (r = .97, p < .001) and not significantly different (t59 = −0.83, p > .40). Reliability was determined using 14 scans of a clinically stable subject acquired at 7 sites, and variability of lesion count was equivalent to that of lesion load. Accounting for lesion load and age, lesion count was negatively associated (t58 = −2.73, p < .01) with the Expanded Disability Status Scale (EDSS). Average lesion size had a higher association with EDSS (r =.35, p < .01) than lesion load (r = .10, p > .40) or lesion count (r = −.12, p > .30) alone.

**Conclusion:** These findings demonstrate that it is possible to recover important aspects of the natural history of lesion formation without longitudinal data, and suggest that lesion size provides complementary information about disease.

**Grant Support:** The project described was supported in part by the NIH grants R01 NS085211, R21 NS093349, and R01 NS094456 from the National Institute of Neurological Disorders and Stroke (NINDS). The study was also supported by the Intramural Research Program of NINDS and the Race to Erase MS Foundation. The content is solely the responsibility of the authors and does not necessarily represent the official views of the funding agencies.

## 1 Introduction

Multiple sclerosis (MS) is a neuroinflammatory disorder characterized by demyelinating lesions that occur in the central nervous system. Magnetic resonance imaging (MRI) is the most commonly used method to observe these lesions, especially in the white matter of the brain^1^. The presence of new lesions on MRI is often considered an important clinical marker of disease activity, yet MRI-based measures of disease severity have been elusive^2^. The total lesion burden in the white matter, or “lesion load” – measured as volume or volume fraction of brain size – is often used in the study of MS, typically as a measure of disease severity^3^ and as a clinical trial outcome^4^. However, lesion load has consistently shown a surprisingly weak association with clinical measures of disease severity, calling into question its usefulness as a surrogate and reinforcing the need for further development of MRI outcomes for MS^2,5^.

In past years, several clinical studies have discussed the number of lesions in a patient’s brain as a possible outcome of interest^6–8^. In these studies, baseline lesion count has been shown to be correlated with EDSS and changes in lesion count have been shown to be correlated with changes in EDSS. However, obtaining an accurate count of biologically distinct lesions in the brain can be costly and logistically challenging, typically requiring expert review of scans taken at regular follow-up visits. This process is especially difficult in patients with a high lesion load and many confluent lesions^9^.

Confluent lesions commonly occur when pathologically distinct lesions (i.e., lesions that arise due to spatially separate sources of structural damage in the brain, usually separated in time) occur in close proximity to each other, creating a larger connected region of lesion tissue. Depending on the level of lesion burden, confluent lesions can range from two overlapping lesions with a single connecting edge to dozens of connected lesions spanning large stretches of white matter. The existence of such confluent tissue can make it difficult or impossible to obtain an accurate estimate of the number distinct lesions in the brain at any given visit. Instead, to determine lesion counts a patient must be scanned regularly, with temporality of appearance serving to separate spatially confluent lesions. However, MRI scans are extremely costly, which can make regular follow-up visits infeasible. Additionally, in patients with a great deal of disease activity, even monthly or bi-monthly scans can produce multiple new lesions that are overlapping in space^10,11^.

To address this issue, the current study introduces a statistical analysis technique for obtaining valid and reliable estimates of lesion count from a single cross-sectional MRI study. This fully automated method utilizes cutting-edge statistical models for segmenting lesion tissue and well-demonstrated mathematical methods for quantifying texture to obtain the number and location of temporally distinct white matter lesions. Additionally, this study provides evidence that the derived lesion counts are associated with clinical measures of disease severity, independent of total lesion volume.

## 2 Methods

### 2.1 Proposed lesion count algorithm

To obtain the lesion count estimate in a given subject, the following steps are carried out.

First, a map of lesion probability at each voxel in the brain is obtained using preprocessed and co-registered MRI volumes from a single visit. Depending on the automated segmentation method that is used, a combination of T_1_-weighted (T1), fluid attenuated inversion recovery (FLAIR), T_2_-weighted (T2), and proton density (PD) volumes will be required for probability estimation. A threshold is then applied to the probability map create a binary mask of regions that are considered lesion tissue.

Using the probability map, the texture of the lesion tissue is quantified to find regions that exhibit the properties expected of the center of a single lesion. Texture is quantified using the eigenvalues of the Hessian matrix. The Hessian matrix is calculated for the intensity of the lesion probability map at every voxel in the lesion mask, with a gradient window of one voxel in each direction. In the context of a 3-dimensional image, the Hessian matrix describes the second-order variation in image intensity in the local neighborhood around a voxel. When applied to a lesion probability map, the eigenvalues of the Hessian matrix at each voxel represent the three primary directions of change in lesion probability at that voxel.

Thus, voxels in the center of a lesion would be expected to have a negative eigenvalue, implying a decrease in probability, in all directions. This follows from the commonly accepted pathology of MS lesions, in which initial damage to a vein causes residual inflammation to spread outwards from the vein in a relatively ovoid fashion, with less damage occurring around the periphery of the visible lesion^12^. Therefore, voxels are eliminated if any of the three eigenvalues are positive, indicating that the voxel is less likely to be lesion than its surroundings in at least one direction. Remaining voxels with three negative eigenvalues are clustered by location, and connected clusters (operationalized as the centers of distinct lesions) are counted.

### 2.2 Data and preprocessing

#### 2.2.1 Validation and clinical-radiological association

Sixty subjects diagnosed with MS were scanned between 2000 and 2008 on a monthly basis over a period of up to 5.5 years (mean = 2.2 years, sd = 1.2) as part of a natural history study at the National Institute of Neurological Disorders and Stroke in Bethesda, Maryland. The subjects ranged from 18 to 60 years of age, with a mean age of 38 years (sd = 9). Of the 60 subjects, 38 were female and 22 were male. The majority of the subjects (n = 44) were diagnosed with relapsing-remitting MS, 13 were characterized as secondary-progressive, one as primary progressive, and two were unspecified. Subjects were either untreated or treated with a variety of disease-modifying therapies during the observation period, including both FDA-approved (various preparations of interferon-beta) and experimental therapies.

Details of the image acquisition and preprocessing have been previously published^13^ and are briefly summarized in this section. Whole-brain 2D FLAIR, PD, T2, and 3D T1-weighted volumes were acquired in a 1.5 tesla (T) MRI scanner (Signa Excite HDxt; GE Healthcare, Milwaukee, Wisconsin). The 2D FLAIR, PD, and T2 volumes were acquired using fast-spin-echo sequences, and the 3D T1 volume was acquired using a gradient-echo sequence. All scanning parameters were clinically optimized for each acquired image. Subjects were each scanned over multiple visits, and subjects’ images at each visit were rigidly co-registered longitudinally and across sequences to a template space^14^.

All images are N4 bias-corrected, and FLAIR, T2, and PD volumes for each subject are interpolated and rigidly co-registered to the T1 volume in isotropic 1 mm^3^ space^15^. Extracerebral voxels were removed using the T1 volume via a skull-stripping procedure^16^, and intensity normalization^17^ of the volumes based on z-scoring was applied. Studies were manually quality controlled by a researcher with over five years’ experience with structural MRI, and studies with analysis-limiting motion or other artifacts were removed. Following preprocessing and quality control, automatic lesion segmentation was performed on co-registered T1, T2, FLAIR, and PD volumes using the OASIS is Automated Statistical Inference for Segmentation (OASIS) model^18^ to produce a lesion probability map for each subject. A conservative threshold of 30% was applied to the probability maps to create binary lesion masks.

#### 2.2.2 Reliability

Data from one 45-year-old man diagnosed with clinically stable relapsing-remitting MS were used to test reliability. This patient was imaged at seven sites in the United States as part of a pilot study for the North American Imaging in Multiple Sclerosis (NAIMS) Cooperative. He was characterized as having mild-to-moderate physical disability, which was stable between the first and last visits, and had no clinical relapses nor radiological changes during the course of the study^19^.

Details of the image acquisition have been previously published^19^ and are briefly summarized in this section. Whole-brain 3D high-resolution FLAIR, T2, and T1-weighted volumes were acquired on seven 3T Siemens MRI scanners across the United States (4 Skyra, 2 Tim Trio, 1 Verio). A standardized high-resolution scanning protocol was developed through a consensus agreement in the NAIMS Cooperative, and was used to the extent possible (allowing for different scanner types and software versions) for each scan. The participant was scanned twice on the same day at each site, and was removed and repositioned between scan and rescan.

All images are N4 bias-corrected, and the subject’s images at each scan were rigidly co-registered across sequences to the T1 volume in isotropic 1 mm^3^ space^15^. Extracerebral voxels were removed using the T1 volume via a skull-stripping procedure^20^, and intensity normalization^17^ of the volumes based on z-scoring was applied. Following preprocessing, automatic lesion segmentation was performed on co-registered T1, T2, and FLAIR volumes using an extension of the OASIS model^21^ to produce a lesion probability map for each scan session. A conservative threshold of 30% was applied to the probability maps to create binary lesion masks.

### 2.3 Statistical analysis

#### 2.3.1 Validation

Using the longitudinal nature of the data, a ‘gold-standard’ count of lesions that appeared during the course of the study was developed for validation. A state-of-the-art technique for segmenting new lesions since a previous visit^22^ was applied at each visit after baseline, resulting in the number and location of new lesions at each visit for every patient. For the gold-standard count, segmented regions containing lesions that were separated in space or time were considered distinct. For example, if a large contiguous region at study’s end consisted of one lesion that appeared at the sixth visit and one lesion that appeared at the eighth visit, this would be considered two lesions in the gold-standard count.

The gold-standard count, henceforth referred to as C_G_, was compared to two counts obtained cross-sectionally at the final of observation for each patient. The first, C_P_, is the count based on the technique proposed in this study. C_P_ was obtained by applying the algorithm described in Section 2.1 to the images obtained at each patient’s final visit, then restricting the count to the number of lesion centers contained in the lesion voxels determined to have appeared during the course of the study. Importantly, this restriction means that C_P_ represents a subset of the total number of lesions in a subject’s scan, and is distinct from the full lesion count that is later described in the context of the clinical-radiological analysis. This limitation was implemented to make direct comparison between C_P_ and C_G_ possible, since a gold-standard count can only be obtained for lesions that appeared during the study.

The second cross-sectional count, C_C_, refers to a count based on the standard connected components technique. C_C_ was obtained by performing lesion segmentation on the images obtained at each patient’s final visit, thresholding at a probability of 30%, and labeling lesions as distinct if they were separated in space. C_C_ was then restricted to the number of unique lesion labels contained in the lesion voxels known to have appeared during the course of the study, in order to facilitate comparison with C_P_ and C_G_.

Comparison between C_G_, C_C_, and C_P_ occurred in two ways. First, to compare the linear correspondence between the gold-standard and the different counting techniques, the correlation between C_G_ and C_P_ was compared to that of C_G_ and C_C_. Then, to determine whether the counts themselves differ meaningfully from the gold-standard, paired t-tests were run for C_G_ and C_P_, as well as C_G_ and C_C_.

#### 2.3.2 Reliability

Determination of the reliability of the proposed counting method was be based on the coefficient of variation (CV) of the counts obtained from the 14 repeated scans. Because the typical connected components technique for counting automatically or manually segmented lesions yields a stable but invalid estimate of the true count, there is no current gold-standard CV for a lesion count. Thus, the CV of the proposed count was compared to a commonly used outcome measure for MS: total cerebral lesion volume (“lesion load”).

This comparison took place in two contexts. The first represented a fully automated version of the proposed count, in which variation may arise from false negatives in the segmentation mask, false positives in the segmentation mask, thresholding of the segmentation mask, and changes in the Hessian structure of the segmentation mask. This coefficient was compared to the CV of automated lesion load, as determined by the segmentation method.

The second context represented a manually supplemented version of the count, where a mask of lesion tissue was provided by an expert rater^19^, and the count was obtained using the segmentation probability map within the manual lesion tissue mask. In this case, variation in the count arises solely due to changes in the Hessian structure of the segmentation mask and changes in the manual segmentation. This coefficient was compared to the CV of the manually obtained lesion load.

#### 2.3.3 Clinical-radiological association

As the Expanded Disability Status Score (EDSS) is known to be noisy, a more stable measure of neurologic disability was created by averaging the EDSS scores over all visits for each subject in the NINDS longitudinal study, hereby referred to as EDSS_avg_. One subject had no EDSS information across all follow-ups, and was excluded from this analysis. Using OASIS lesion probability maps^18^, lesion load was obtained at the final visit for each subject using a probability threshold of 30%. Then, using the lesion count technique described in Section 2.1, a full count of white matter lesions at the final visit was obtained for each subject. Importantly, the counts obtained for the clinical-radiological analysis are distinct from the C_P_ measure described in Section 2.3.1, as these counts represent the application of the proposed method to the entire brain, while C_P_ represents the application of the proposed method to only lesion tissue that appeared during the course of the longitudinal study.

To determine the clinical relevance of the proposed lesion count independent of other potentially confounding variables, a linear regression model was created for EDSS_avg_ with age, lesion load, and lesion count as predictors. Additionally, Pearson correlations with EDSS_avg_ were calculated for lesion load and lesion count, as well as a new variable we refer to as average lesion size (defined as lesion load divided by lesion count).

## 3 Results

### 3.1 Validation

The temporally informed gold-standard count of new lesions appearing over the course of study, C_G_, ranged from 0 to 75 among the 60 subjects, with a median of 4 (*IQR* = [1, 12]). The connected components count, C_C_, ranged from 0 to 14 with a median of 2 (*IQR* = [1, 5]). The proposed count, C_P_, ranged from 0 to 60 with a median of 5 (*IQR* = [1, 15]). Figure 1 provides a visual example of these counting techniques.

**Figure 1.**
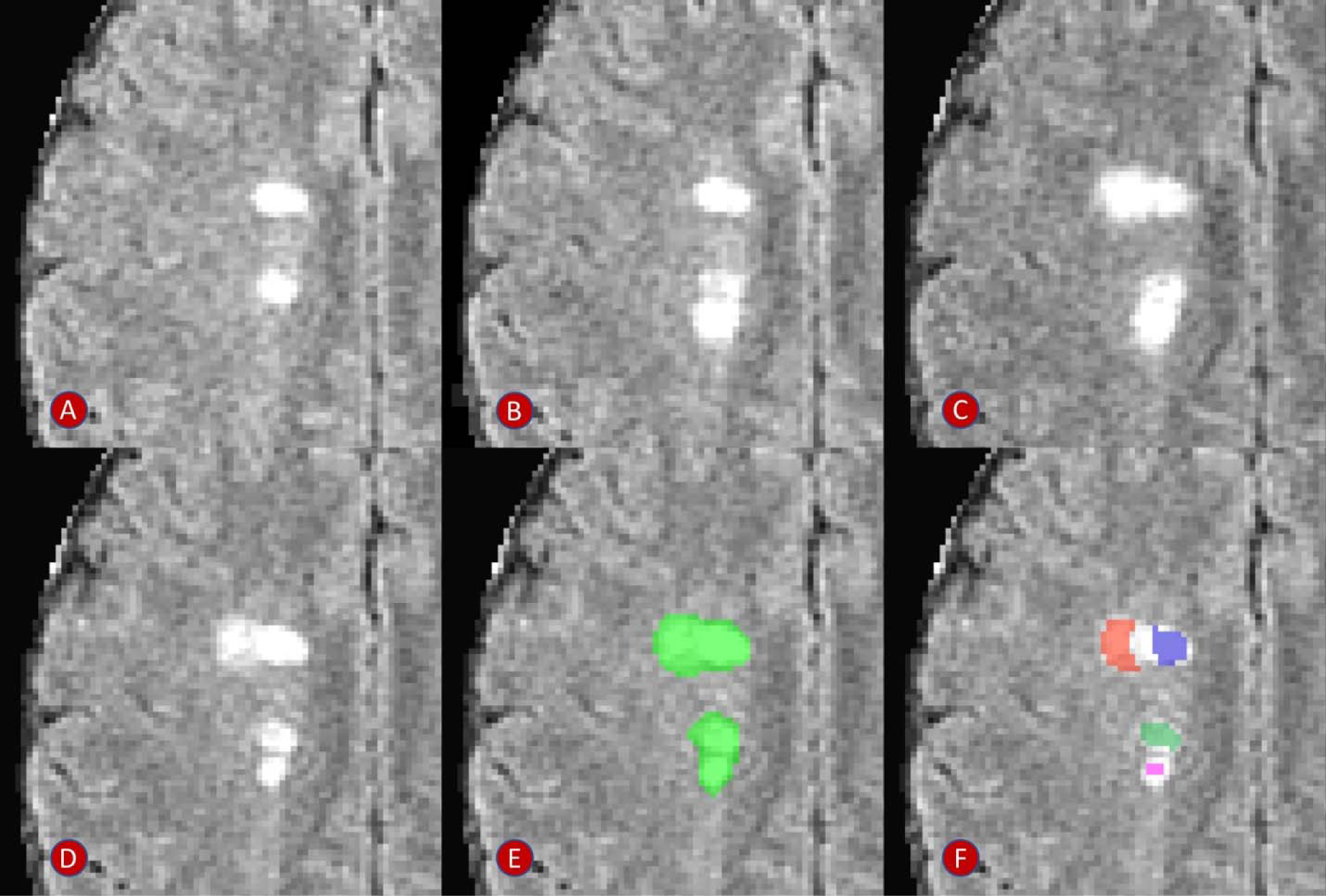
Example of the lesion counts in a region with four apparently distinct lesions, two of which develop with observable temporal separation. Panels A-D show development of two new and temporally distinct lesions. Panels E and F show the performance of a connected components count and the proposed count, respectively. The connected components method finds one confluent lesion in the visualized space (connected in an adjacent plane), and the proposed method finds four distinct lesion centers. Days from scan in panel A: (B) 28 days; (C) 91 days; (D-F) 252 days.

The correlation between C_P_ and C_G_ was .97, compared to the correlation of .67 between C_C_ and C_G_. Figure 2 shows the scatterplots for the two linear associations, along with the line demonstrating a one-to-one relationship. The paired t-test comparing C_C_ and C_G_ yielded a highly significant result (*t*_59_ = 4.19, *p* < .001), with C_G_ being 6.9 lesions larger than C_C_ on average (95% CI: [3.6, 10.2]). The paired t-test comparing C_P_ and C_G_ did not find a significant difference between the counts (*t*_59_ = −0.83, *p* > .40), with C_P_ being 0.4 lesions larger than C_G_ on average (95% CI: [−1.3, 0.5]).

**Figure 2.**
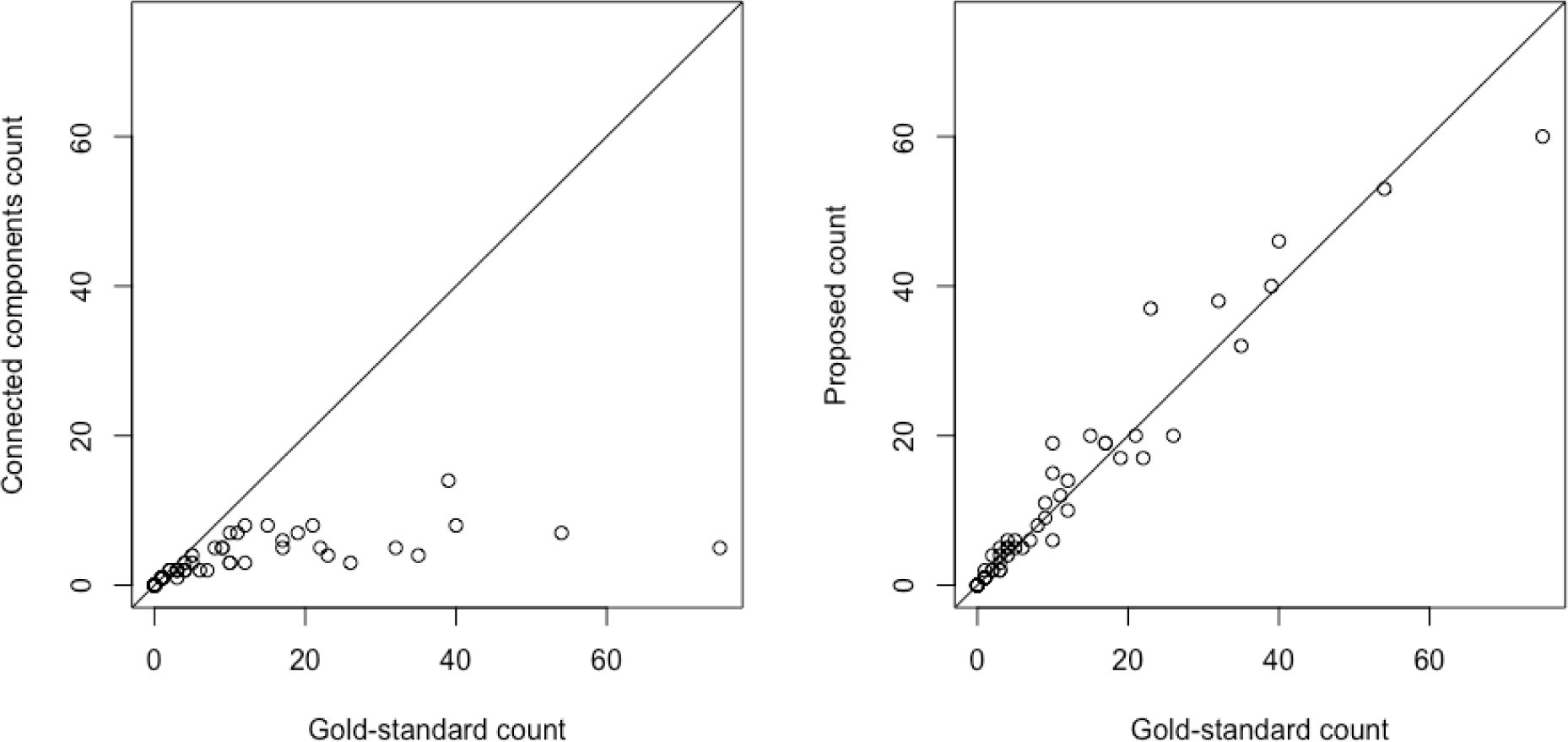
Scatterplots for the comparison between the gold-standard count and the connected components count and comparison between the gold-standard count and the proposed count, respectively. Diagonal lines represent a one-to-one relationship.

### 3.2 Reliability

For the fully automated count, the coefficient of variation was .19, compared to a CV of .22 for the automated lesion load. Using the manual segmentation as a mask, the CV for the lesion count was reduced to .12, compared to a CV of .10 for the manual lesion load. In one case, automated lesion segmentation was discovered to have failed, creating a probability map with a drastically different Hessian structure and large regions of false positive segmentation. With this scan removed the CV of the fully automated lesion count remained at .19 and the CV of the manual segmentation-based lesion count dropped to less than .06, suggesting that the proposed count has equivalent or lower variability than the current clinical standard of lesion load.

### 3.3 Clinical-radiological association

Accounting for lesion load and age, the proposed lesion count was negatively associated with EDSS_avg_ (t_58_ = −2.73, p < .01), suggesting that for a given lesion load and age, a higher count is associated with lower disease severity. The inclusion of lesion count in the model explains an additional 10% of the variance in EDSS_avg_ compared to a model with only age and lesion load, providing support to the hypothesis that the proposed count contains disease information independent of other commonly used measures.

The Pearson correlation between lesion load and EDSS_avg_ was small and did not reach significance (*r* = .10, *p* > .40), nor did the correlation between lesion count and EDSS_avg_ (*r* = − .12, *p* > .30). However, average lesion size was significantly correlated with EDSS_avg_ (*r* = .35, *p* < .01), indicating that larger lesions were associated with higher disability.

## 4 Discussion

In this paper, we introduce a novel technique for obtaining cross-sectional counts of pathologically distinct lesions, and demonstrate it to be a valid, reliable, and clinically meaningful biomarker for MS disease status. Utilizing information contained in the Hessian structure of lesion probability maps produced by automated segmentation methods, this technique counts distinct lesions by identifying regions that resemble the physiological traits of distinct lesion centers.

Validity of this measure was established by comparing counts obtained at a single time point to gold-standard counts that incorporated temporal information on lesion development. The proposed count had a correlation of .97 with the gold-standard count, indicating very strong validity of this measure. A count obtained using the connected components method had only a .67 correlation with the gold-standard, and appeared to strongly underestimate the number of lesions in individuals who developed more than one or two lesions per year over the course of the study. This underestimation manifested in a highly significant difference between the connected components counts and the gold-standard counts in a paired t-test, whereas no difference was found between the proposed counts and the gold-standard counts. These findings demonstrate that the proposed technique yields a count that is consistent with the natural history of lesion formation.

Reliability was considered using a rich set of data from the NAIMS Cooperative. In that study, a clinically and radiologically stable subject was scanned two times at each of seven different sites across the United States. To judge the reliability of the proposed measure, the lesion count was obtained for all 14 scans of this subject, and the coefficient of variation of the counts was compared to that of lesion load in two contexts. In the fully automated comparison, lesion count had a slightly lower CV than lesion load. This indicates that across repeated scans of the same brain, automated lesion count is a less variable measure than automated lesion load. In the manually supplemented comparison, lesion count had a slightly higher CV than lesion load, implying that manually obtained lesion load is a slightly less variable measure than semi-automated lesion count. Upon inspection there appeared to be one scan where automated lesion segmentation failed, producing an abnormal Hessian structure within the manually segmented lesion mask. With this scan removed, the CV of semi-automated lesion count dropped to slightly more than half that of manual lesion load. This suggests that when automated lesion segmentation methods perform as expected, semi-automated lesion count is appreciably more reliable than manual lesion load, a widely used measure of disease severity.

Clinically, the lesion count measure appears to be a potentially important addition to commonly used radiological biomarkers for MS. In a model accounting for lesion load and age, lesion count was highly significantly associated with EDSS. Interestingly, this association was negative, indicating that for subjects who have similar lesion load, better outcomes are associated with more (and smaller) lesions rather than fewer (and larger) lesions. This lends support to the idea that neither the number of lesions nor the amount of tissue damage alone captures all relevant clinical information, and instead that suggests they should be considered together. One way to conceptualize the combination of these metrics is average lesion size, which taps into the degree to which the brain is capable of halting the growth of lesions and encouraging lesional recovery^13,23,24^ after incidence.

To investigate this concept more directly, a measure of average lesion size was created by dividing lesion load by lesion count. Pearson correlations with EDSS were then compared for the three biomarkers of lesion load, lesion count, and average lesion size. These findings provided further support for the combined importance of lesion load and lesion count, with both showing small and nonsignificant associations with EDSS. However, average lesion size showed a significant positive association with EDSS, consistent with the notion that the brain’s ability to slow or stop lesion growth is clinically relevant. These findings point to the importance of considering lesion count in MS research, and provide further evidence of the validity of the proposed counting technique.

The main limitation of the current study is the possibility of alternate explanations of confluence that are not accounted for in the design of the proposed count. It has been hypothesized that confluent lesions may occasionally occur as a result of the growth of older lesions, or the expansion of pathological processes. Future research should consider the degree to which this technique does or does not characterize these types of confluence as pathologically distinct lesions.

The lesion count method presented in this paper has several appealing features, including its low computational burden and its easy and flexible implementation. Computationally, the counting algorithm takes less than a minute to run once probability maps are obtained. The speed of the full technique varies depending on the lesion segmentation method used, but took approximately 25 minutes per subject as presented in this study. In terms of implementation, this method can be quickly and easily coded in any program capable of calculating the Hessian structure of a 3D image, a feature included in most image processing packages. It can also be used with any lesion segmentation method that yields a probability map, allowing it be added to almost any pipeline regardless of preferred segmentation algorithm.

## 5 Conclusion

This paper introduces a novel and reliable fully automated method for counting pathologically distinct lesions using images obtained at a single time point, allowing for an accurate reconstruction of the natural history of lesion formation without longitudinal data. Lesion count was found to be significantly associated with EDSS, independent of potential confounders such as lesion load and age, and the results suggest that individuals with more small lesions may have better clinical outcomes than those with fewer large lesions. This study also demonstrates the importance of obtaining both lesion count and lesion load by using them to construct a new MS biomarker, average lesion size, and showing that average lesion size has a significantly larger association with EDSS than both lesion load and lesion count. With further study, this technique and the findings it produces could set the stage for new lesion-level considerations in research and treatment of MS.

## 6 Acknowledgments

The authors would like to thank Blake Dewey, the Neuroimmunology Branch clinical group and the technicians at the NIH who were instrumental in helping to collect and process the validation and clinical-radiological data. Additionally, the following is a full list of individuals who contributed to the collection and processing of the NAIMS data used in the reliability analysis described in this paper: Brigham and Women’s Hospital, Harvard Medical School (Boston, MA): Rohit Bakshi, Renxin Chu, Gloria Kim, Shahamat Tauhid, Subhash Tummala, Fawad Yousuf; Cedars-Sinai Medical Center (Los Angeles, CA): Nancy L. Sicotte; Henry M. Jackson Foundation for the Advancement of Military Medicine (Bethesda, MD): Dzung Pham, Snehashis Roy; National Institutes of Health (Bethesda, MA): Frances Andrada, Irene C.M. Cortese, Jenifer Dwyer, Rosalind Hayden, Haneefa Muhammad, Govind Nair, Joan Ohayon, Daniel S. Reich, Pascal Sati, Chevaz Thomas; Johns Hopkins (Baltimore, MD): Peter A. Calabresi, Sandra Cassard, Jiwon Oh; Oregon Health and Science University (Portland, OR): William Rooney, Daniel Schwartz, Ian Tagge; University of California (San Francisco, CA): Roland G. Henry, Nico Papinutto, William Stern, Alyssa Zhu; University of Pennsylvania (Philadelphia, PA): Christos Davatzikos, Jimit Doshi, Guray Erus, Kristin Linn, Russell Shinohara; University of Toronto (Ontario, Canada): Jiwon Oh; Yale University (New Haven, CT): R. Todd Constable, Daniel Pelletier.

## References

1. MRI Atlas of MS Lesions. (Springer Berlin Heidelberg, 2008).

2. Barkhof, F. The clinico-radiological paradox in multiple sclerosis revisited. Curr. Opin. Neurol. 15, 239–245 (2002).

3. Popescu, V. et al. Brain atrophy and lesion load predict long term disability in multiple sclerosis. J. Neurol. Neurosurg. Psychiatry 84, 1082–1091 (2013).

4. Calabresi, P. A. et al. Safety and efficacy of fingolimod in patients with relapsing-remitting multiple sclerosis (FREEDOMS II): a double-blind, randomised, placebo-controlled, phase 3 trial. Lancet Neurol. 13, 545–556 (2014).

5. Thompson, A. J. et al. Patterns of disease activity in multiple sclerosis: clinical and magnetic resonance imaging study. BMJ 300, 631–634 (1990).

6. Brex, P. A. et al. A Longitudinal Study of Abnormalities on MRI and Disability from Multiple Sclerosis. N. Engl. J. Med. 346, 158–164 (2002).

7. Khoury, S. J. et al. Longitudinal MRI in multiple sclerosis: correlation between disability and lesion burden. Neurology 44, 2120–2124 (1994).

8. Rudick, R. A., Lee, J.-C., Simon, J. & Fisher, E. Significance of T2 lesions in multiple sclerosis: A 13-year longitudinal study. Ann. Neurol. 60, 236–242 (2006).

9. Zivadinov, R., Zorzon, M., De Masi, R., Nasuelli, D. & Cazzato, G. Effect of intravenous methylprednisolone on the number, size and confluence of plaques in relapsing–remitting multiple sclerosis. J. Neurol. Sci. 267, 28–35 (2008).

10. Harris, J. O., Frank, J. A., Patronas, N., McFarlin, D. E. & McFarland, H. F. Serial gadolinium-enhanced magnetic resonance imaging scans in patients with early, relapsing-remitting multiple sclerosis: Implications for clinical trials and natural history. Ann. Neurol. 29, 548–555 (1991).

11. Guttmann, C. R. et al. Multiple sclerosis lesion formation and early evolution revisited: A weekly high-resolution magnetic resonance imaging study. Mult. Scler. J. 22, 761–769 (2016).

12. Gaitán, M. I. et al. Evolution of the blood-brain barrier in newly forming multiple sclerosis lesions. Ann. Neurol. 70, 22–29 (2011).

13. Sweeney, E. M. et al. Relating multi-sequence longitudinal intensity profiles and clinical covariates in incident multiple sclerosis lesions. NeuroImage Clin. 10, 1–17 (2016).

14. Fonov, V. et al. Unbiased average age-appropriate atlases for pediatric studies. NeuroImage 54, 313–327 (2011).

15. Avants, B., Tustison, N., Song, G. & Gee, J. ANTS: Advanced Open-Source Normalization Tools for Neuroanatomy. (Penn Image Computing and Science Laboratory, 2009).

16. Carass, A. et al. A joint registration and segmentation approach to skull stripping. in Biomedical Imaging: From Nano to Macro, 2007. ISBI 2007. 4th IEEE International Symposium on 656–659 (IEEE, 2007).

17. Shinohara, R. T. et al. Statistical normalization techniques for magnetic resonance imaging. NeuroImage Clin. 6, 9–19 (2014).

18. Sweeney, E. M. et al. OASIS is Automated Statistical Inference for Segmentation, with applications to multiple sclerosis lesion segmentation in MRI. NeuroImage Clin. 2, 402–413 (2013).

19. Shinohara, R. T. et al. Volumetric Analysis from a Harmonized Multisite Brain MRI Study of a Single Subject with Multiple Sclerosis. Am. J. Neuroradiol. 38, 1501–1509 (2017).

20. Jenkinson, M., Pechaud, M. & Smith, S. BET2: MR-based estimation of brain, skul and scalp surfaces. (2005).

21. Valcarcel, A. et al. MIMoSA: A Method for Inter-Modal Segmentation Analysis. (2017). doi:10.1101/150284

22. Sweeney, E. M., Shinohara, R. T., Shea, C. D., Reich, D. S. & Crainiceanu, C. M. Automatic Lesion Incidence Estimation and Detection in Multiple Sclerosis Using Multisequence Longitudinal MRI. Am. J. Neuroradiol. 34, 68–73 (2013).

23. Meier, D. S. & Guttmann, C. R. G. Time-series analysis of MRI intensity patterns in multiple sclerosis. NeuroImage 20, 1193–1209 (2003).

24. Meier, D. S. & Guttmann, C. R. G. MRI time series modeling of MS lesion development. NeuroImage 32, 531–537 (2006).

